# Nicking Loop™: Streamlined circular DNA libraries for precision genomics, DNA data storage and universal NGS read-out

**DOI:** 10.1101/2025.05.08.652165

**Authors:** Simona Adamusová, Nea Laine, Anttoni Korkiakoski, Tatu Hirvonen, Anna Musku, Tuula Rantasalo, Jorma Kim, Juuso Blomster, Jukka Laine, Manu Tamminen, Juha-Pekka Pursiheimo

## Abstract

Circular single-stranded DNA (CssDNA) offers unique advantages for molecular diagnostics and next-generation sequencing (NGS) due to its nuclease resistance and compatibility with rolling circle amplification. We present the Nicking Loop™, a novel and versatile method for converting both single- and double-stranded DNA into CssDNA and enabling robust, unbiased amplification. This technique preserves the original DNA template composition and outperforms PCR in reproducibility and sensitivity, particularly for low variant allele frequencies. We demonstrate that Nicking Loop-amplified CssDNA is directly compatible with NGS platforms utilizing circular DNA as a sequencing template and introduce a novel loop-based indexing strategy that enables efficient sample multiplexing. Furthermore, we confirm the method’s compatibility with nanopore sequencing technologies, highlighting its broad applicability across sequencing platforms. Our results establish Nicking Loop™ as an alternative to conventional linear library preparation, streamlining workflows while enhancing sequencing accuracy and efficiency. Beyond its immediate application in NGS, the method holds promise for broader use in molecular biology, e.g. in DNA data storage. This proof-of-concept study highlights the transformative potential of Nicking Loop™ in enabling a shift toward circular DNA libraries for new generation of assay preps and precision sequencing applications.

## INTRODUCTION

Circular single-stranded DNA (CssDNA) is emerging as a cornerstone in modern biotechnology (1), with growing significance in next-generation sequencing (NGS) applications (2–7). The covalently closed structure of the CssDNA offers unique benefits over the linear DNA. For instance, the CssDNA is inherently resistant to exonuclease degradation, granting a superior biostability under various conditions. Additionally, CssDNA serves as an ideal template for rolling circle amplification (RCA), enabling robust linear amplification making it valuable for a broad range of applications, e.g. in diagnostics or DNA data storage.

RCA is an isothermal nucleic acid amplification method employing a strand-displacing polymerase to generate long concatemeric single-stranded DNA (ssDNA) products consisting of tandem repeats of the original template. RCA amplifies DNA linearly, resulting in moderate amounts of product DNA that is adequate for several applications but may be insufficient for those requiring higher level of DNA amplification (8). Therefore, to broaden the use of RCA and circular templates, efficient amplification strategies permitting direct multiplication of circular templates are needed. A well-known approach to improve template availability and enhance the yield of RCA is circle-to-circle amplification (9). In this method, a strand-specific amplification of CssDNA is performed in cycles, where RCA-derived concatemers are enzymatically cleaved into monomeric units, which are then re-circularized to serve as templates for subsequent RCA. However, each step requires an addition of specific splinting oligonucleotides, introducing methodological complexity and potential reduction of reaction efficiency (10).

This study presents the Nicking Loop™, a novel technique for converting both ssDNA and double-stranded DNA (dsDNA) into CssDNA, enabling robust and direct amplification of CssDNA molecules. The core components of the system include Nicking Loop™ oligonucleotide (hereafter referred as Loop), bridge oligonucleotide and target-specific probe pair. These oligonucleotides form a complex in which Loop and probes are stabilized by the bridge oligonucleotide. Upon hybridization to target DNA, the construct is circularised using a combined action of a DNA polymerase and ligase.

The Loop is a fundamental part of the Nicking Loop™ method. After RCA, the repeated Loop structure within the long concatemeric molecule folds into a hairpin form that facilitates site-specific nicking and cleavage of the concatemer into monomeric units. These monomeric units undergo Loop-mediated self-folding followed by ligation to form circular structures. This eliminates the need for constant addition of external splinting oligonucleotides, enhancing reaction efficiency and drastically simplifying the workflow. In addition to enabling circularization and amplification, loop-region of the Loop can be engineered to incorporate unique molecular identifiers (UMIs) as well as sample-specific indices, allowing the efficient multiplexing and early sample pooling. This permits early indexing of the samples within the workflow and significantly improves cost-efficiency of the process by facilitating parallel sequencing of multiple, separately indexed samples. Importantly, this method enables direct sequencing of CssDNA by incorporating sequencing adapter motifs into the circular molecules allowing fast and easy library preparation. Taken together, these features pave the way for entirely new assay preparation paradigm, transitioning from conventional linear library structures to more versatile and robust circular library preparation and sequencing strategies.

## MATERIAL AND METHODS

### Oligonucleotides

The basic components of the Nicking Loop™ include: (i) a bridge oligonucleotide that binds to (ii) left and right probes, (iii) the Loop that hybridizes to bridge oligonucleotide and completes the gap between the probes, and (iv) targets captured by the probes. Two synthetic target pools, ten probes, bridge and Loops were produced by IDT (Coralville, IA). The synthetic target pools comprise ten shared templates, with each pool distinguished by three pool-specific complementary nucleotides. The pools were mixed in different ratios to mimic variant allele frequency (VAF) of 0%, 1%, 5%, 10% and 20%.

### Nicking Loop™ conversion of linear DNA to CssDNA

To convert linear DNA to CssDNA, 2.5 fmol of synthetic target pools were combined with 10 fmol of probes (two specific probes loaded at 1 fmol to balance performance) and 200 fmol of Loop in 1× Ampligase buffer (LGC Biosearch Technologies, Hoddesdon, UK). To induce hybridization of the components, the mixture was denatured at 95°C for 5 minutes and temperature was gradually decreased (10°C step, hold 2.5 minutes) to 55°C and incubated for 2 hours. The hybridized components were circularized with 0.2 U of Phusion High-Fidelity DNA Polymerase (Thermo Fisher Scientific, Waltham, MA), 0.2 U Ampligase (LGC Biosearch Technologies), and 10 mM dNTP (Thermo Fisher Scientific) in 1× Ampligase Reaction buffer (LGC Biosearch Technologies) and incubated at 55°C for 40 minutes. To remove residual linear DNA, 1 µL each of Thermolabile Exonuclease I, RecJf and Lambda Exonuclease (NEB, Ipswich, MA) was added to the reaction and incubated at 37°C for 1 hour, followed by heat inactivation at 80°C for 10 minutes. The reaction was then purified using a 2× volume of AMPure XP beads (Beckman Coulter, Brea, CA). At this stage, linear DNA was transformed into Nicking Loop™-converted CssDNA, enabling subsequent amplification.

### PCR control for Nicking Loop™ conversion

To assess the efficiency of the conversion, 2 µL of Nicking Loop™-converted CssDNA was subjected to PCR. The primers were designed to anneal to regions flanking the Loop insertion site, enabling the amplification only if the DNA molecule had been successfully circularized. The reaction was supplemented with 10 pM of primers (IDT), 0.02 U of Phusion HS II DNA Polymerase (Thermo Fisher Scientific), and 10 mM dNTP (Thermo Fisher Scientific). The thermocycler program was based on manufacturer’s guidelines with annealing temperature of 68°C and 12 cycles. The gel electrophoresis and DNA quantification was performed using High Sensitivity D1000 ScreenTape and reagents with 4150 TapeStation System (Agilent, Santa Clara, CA).

### Nicking Loop™ amplification

To amplify the circularized molecules, 10 µL of Nicking Loop™-converted CssDNA was pre-annealed with 5 pmol of universal amplification primer in 1× EquiPhi29 buffer (Thermo Fisher Scientific). The mixture was incubated at 95°C for 3 minutes and then gradually cooled to room temperature to facilitate primer annealing to the CssDNA template. RCA was initiated by adding 10 mM dNTPs, 100 mM DTT and 0.5 U of EquiPhi29 (Thermo Fisher Scientific) followed by incubation at 45°C for 90 minutes. The reaction was terminated by heat inactivation of the enzyme at 95°C for 10 minutes.

The newly synthetized concatemeric ssDNA was supplemented with nuclease-free water and 1× rCutSmart buffer (NEB). The supplemented reactions were heated to 95°C for 5 minutes and then gradually cooled to room temperature to promote Loop folding. The double-stranded segment of the folded Loops contained recognition site for nicking enzyme. The recognition site was digested with 10 U of Nb.BsrDI (NEB) at 37°C for 90 minutes resulting into linear monomers. The enzyme was subsequently heat-inactivated at 80°C for 10 minutes. The resulting linear monomers were purified using 1.8× volume of AMPure XP beads (Beckman Coulter).

To initiate re-circularization, 10 µL of linear monomers were mixed with 7 µL of nuclease-free water. The mixture was denatured at 95°C for 5 minutes and gradually cooled to room temperature to allow folding of Loop. Subsequently, 1× T4 DNA ligase buffer and 1 µL of T4 DNA ligase (NEB) were added to each reaction to ligate the nick in the folded monomer. Ligation was carried out at 32°C for 30 minutes. After the ligation, any residual ssDNA was removed by adding 1 µL each of Thermolabile Exonuclease I and RecJf (NEB). The reactions were incubated at 37°C for 45 minutes, then heat-inactivated at 95°C for 10 minutes. The reactions were purified using 2× volume of AMPure XP beads (Beckman Coulter). After this step, the original CssDNA molecules had been successfully amplified resulting in a first-generation of amplified molecules. This entire process RCA, Loop digestion, and re-circularization – was repeated up to three times using the newly generated circular molecules as templates and a reverse amplification primer to produce subsequent generations of CssDNA molecules.

### Quantification of molecules produced in Nicking Loop™ amplification

The number of molecules produced in each generation of Nicking Loop™ amplification was quantified using quantitative real-time PCR (qPCR). The reaction consisted of 1 µL of template, 2.5 µM of proprietary primers and 5 µL of SsoAdvanced Universal SYBR Green Supermix (Bio-Rad, Hercules, CA), and was filled with nuclease-free water to final volume of 10 µL. Either linear molecules originating from monomerization step of Nicking Loop™ amplification, synthetic standards (IDT), or nuclease-free water (as a negative control) were used as the template. The qPCR reaction was performed using a Biorad CFX96 Touch Real-Time PCR System (Bio-Rad). The thermal cycle was programmed for 2 minutes at 98°C for initial denaturation, followed by 27 cycles of 15 seconds at 98°C for denaturation and 45 seconds at 67°C for annealing and extension.

The optimal reaction conditions were determined empirically. Optimal annealing temperature was selected from range of 54°C–72°C, and primer concentration was selected from range of 0.5 µM–10 µM. Standard curve was prepared in eight 2-fold dilutions of synthetic standard with 5 technical replicates at each dilution. To quantify linear molecules, three 2-fold dilutions with five technical replicates were prepared. The cycle threshold values were determined using CFX Maestro software version 4.1 (Bio-Rad). The slope of the calibration curve was −3.42, with a *y* intercept of 42.56, R^2^ value of 0.98 and reaction efficiency of 95.95%.

### PCR amplified generations

To compare the Nicking Loop™ amplification with conventional PCR, 1 µL of the first-generation Nicking Loop™ amplified libraries were amplified by PCR for 15 cycles, generating the first PCR-derived generation. Subsequent generations were produced in the same manner, with each PCR-generation serving as the template for the next round of amplification. All PCR products were gel-purified using Monarch DNA Gel Extraction kit (NEB). Purified libraries were quantified using Qubit™ dsDNA HS Assay Kit and Qubit™ 4 Fluorometer (Thermo Fisher Scientific). Normalized libraries were sequenced on Illumina MiSeq platform, using the MiSeq Reagent Kit v3 for a 2 × 300 bp paired-end run (Illumina, San Diego, CA).

### Loop incorporation efficiency

To assess whether structural differences in the Loop affect its incorporation efficiency into Nicking Loop™-converted structure, a competitive incorporation assay was performed. Twenty-five distinct Loops were mixed in an equimolar ratio to create a pool, each containing a different central nucleotide sequence within the loop-region (Supplementary Table S4). A total of 200 fmol of pool was mixed with 2.5 fmol of synthetic target and 10 fmol of corresponding probe and supplemented with 1× Ampligase buffer (LGC Biosearch Technologies). The subsequent Nicking Loop™ conversion and circularization steps were performed as described in Nicking Loop™ conversion of linear DNA to circular DNA of methods section. The resulting products were amplified by Nicking Loop™ up to first-generation (see Nicking Loop™ amplification). Final libraries were sequenced on Illumina iSeq instrument using iSeq 100 i1 Reagent v2 (300-cycle) (Illumina). All reactions were performed in 10 independent replicates.

### Library preparation and sequencing with Illumina MiSeq

For sequencing by Illumina, 2 µL of CssDNA molecules from each of the generations were subjected to PCR using 10 µM of dual-indexed primers with Phusion HS II DNA Polymerase (Thermo Fisher Scientific) following the manufacturer’s guidelines. PCR was performed with annealing temperature of 68°C for 15 cycles. All PCR products were gel-purified using the Monarch DNA Gel Extraction kit (NEB). The libraries were quantified using Qubit™ dsDNA HS Assay Kit and Qubit™ 4 Fluorometer (Thermo Fisher Scientific). Sequencing was performed on Illumina MiSeq platform, using the MiSeq Reagent Kit v3 for a 2 × 300 bp paired-end run (Illumina).

### Library preparation and direct sequencing of CssDNA libraries

To evaluate direct compatibility with NGS platforms that utilize CssDNA in their workflows, Nicking Loop™ components were designed to incorporate all required functional motifs from the outset. Nicking Loop™ conversion and Nicking Loop™ amplification reactions were performed as previously described. CssDNA libraries were purified using 1.8× volume of AMPure XP beads (Beckman Coulter) and quantitated using Qubit™ ssDNA HS Assay Kit and Qubit™ 4 Fluorometer (Thermo Fisher Scientific). Libraries were sequenced with a modified protocol allowing direct analysis of CssDNA libraries.

### Library preparation and sequencing with Oxford Nanopore

To evaluate Nicking Loop™ compatibility with the Oxford Nanopore sequencing, the amplified RCA concatemers were prepared from Nicking Loop™-converted circles as previously described. The concatemers were purified with the 1× AMPure XP Beads (Beckman Coulter) and quantified using Qubit™ ssDNA HS Assay Kit and Qubit™ 4 Fluorometer (Thermo Fisher Scientific). The samples were barcoded with Rapid Barcoding Kit V14 (Oxford Nanopore Technologies, Oxford, UK) and sequenced using MinION with MinION or GridION Flow Cell v.R10.4.1 (Oxford Nanopore technologies).

### Data analysis

Reads generated by Illumina sequencers were merged using VSEARCH (11) (v2.15.2_linux_x86_64) with the following parameters: --fastq_minovlen 10 --fastq_maxdiffs 15 --fastq_maxee 1 --fastq_allowmergestagger --fastq_qmaxout 92. For both Nicking Loop™ and PCR amplified generations, the original forward and reverse reads were subsampled without replacement to 97,857 reads prior to merging, and only merged reads with a quality score of 60 or higher were retained for further analysis.

Otherwise, all data were processed using proprietary pipelines. Two probes, which had been diluted tenfold relative to the rest of the panel, were excluded from the Nanopore sequencing data analysis due to insufficient sequencing depth. Python version 3.11.5 and matplotlib library version 3.7.2 were used to draw the figures. SciPy (v1.11.1) library was used for Spearman correlations.

## RESULTS

### Nicking Loop™-converted DNA is compatible with linear amplification cycles

Nicking Loop™ method provides a unique approach to convert a linear DNA to a CssDNA form (Fig. 1A) enabling efficient linear amplification (Fig. 1B). The resulting circular molecules can be repeatedly amplified, with each cycle producing new generation of the original circle with alternating strand polarity.

**Figure 1.**
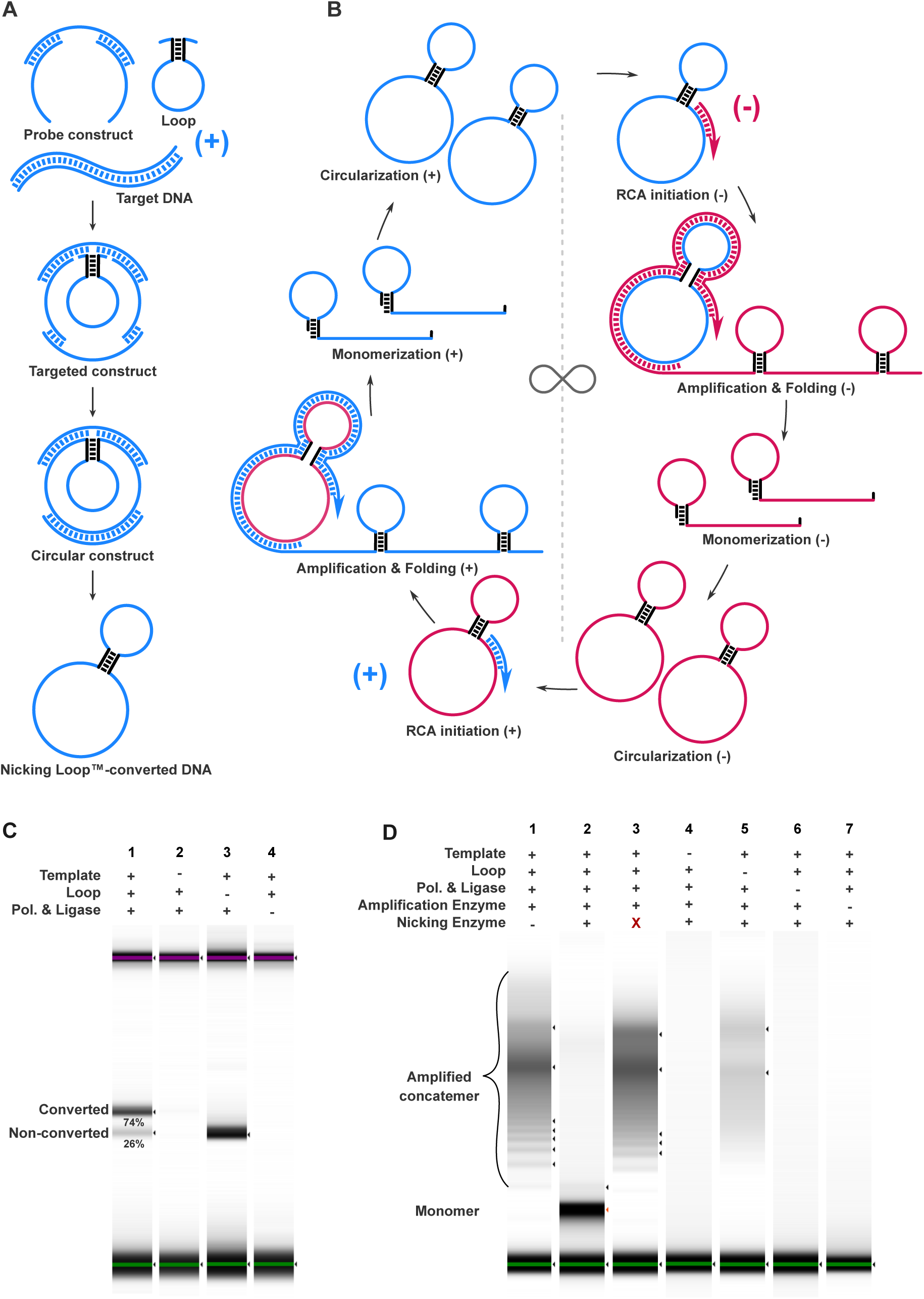
Process of Nicking Loop™ conversion and amplification. **(A)** Nicking Loop™ conversion starts with addition of the probe constructs and Loop to linear target DNA. The Loop forms the hairpin structure with a restriction site located at the complementary stem (black). The overhangs of the Loop binds to the bridge in the gap between the probes, and the target specific probe arms capture the target DNA, forming a targeted construct. The gap of the targeted construct is filled by polymerase and the nicks in the structure are ligated, resulting in the circular construct. Any residual linear DNA is degraded by exonucleases leaving intact Nicking Loop™-converted circles. **(B)** Nicking Loop™ amplification begins with priming of Nicking Loop™-converted circle, leading to RCA initiation. The circle is amplified into a single-stranded DNA (ssDNA) concatemer, which folds into distinct Loop structure with restriction site in the stem. During monomerization, the stem of the Loops is cleaved with a nicking enzyme, breaking concatemer into monomers. The monomers are subsequently folded into a circular structure with a nick in the hairpin stem, which is then ligated to complete the circularization, and any residual linear DNA is degraded by exonucleases. This process can be infinitely repeated, each time creating a new generation with opposite strand polarity. The (+) and (-) indicates the changing polarity of the strands. **(C)** Nicking Loop™ conversion demonstrated by gel electrophoresis (Tapestation – HS DNA ScreenTape). The Nicking Loop™ conversion efficiency was reflected by a high abundance of Nicking Loop™-converted circles compared to non-converted products [1]. The non-converted circular products were formed in absence of the Loop [3]. In the absence of template [2] or GapFill enzymes [4] no product was formed. **(D)** The monomerization step in the first generation of Nicking Loop™ analyzed by gel electrophoresis (Tapestation – HS RNA ScreenTape). In the absence of the nicking enzyme [1] or if nicking enzyme is incompatible with recognition site (X) [3], the amplified concatemer remained intact. In presence of compatible nicking enzyme [2], the concatemer was digested into the monomers. No product was formed in absence of template [4], GapFill enzymes [5] or amplification enzyme [7] during Nicking Loop™ conversion. Without the Loop, a non-converted circular molecule was able to produce long concatemeric molecule but failed to monomerized and was degraded by exonucleases before completion of first amplification cycle.

Conversion of linear DNA into CssDNA by the Nicking Loop™ requires the following components: the Loop, a bridge oligonucleotide, a pair of target-specific probe oligonucleotides, and the linear target DNA. The Loop forms a hairpin structure with overhangs, a complementary stem carrying a nicking endonuclease recognition site and a loop-region that contains the functional motifs for sample indexing and/or UMIs. The bridge oligonucleotide anneals with the left and right probes, forming a probe construct and leaving a central gap for binding of the Loop overhangs. The probe arms contain target specific sites which bind to the target DNA, creating a gap between the probes and forming a targeted construct. This gap is consequently filled by polymerase, copying the target DNA. The copied target DNA, the probes and the Loop are ligated together, creating a circular construct. To complete the conversion, the bridge oligonucleotide and any residual linear DNA are degraded by exonucleases, resulting in intact Nicking Loop™-converted CssDNA molecules (Fig. 1A).

The Nicking Loop™-converted circles can be linearly amplified through repeated cycles of RCA, Loop folding, nicking-enzyme mediated monomerization, and circularization – each cycle producing a new generation of circles with alternating strand polarity. The RCA is initiated from the universal amplification primer pre-annealed to the circular molecule, resulting in ssDNA concatemer. Repeats of the newly synthetized concatemer fold into distinct Loop structure to form a nicking endonuclease recognition site. In the next step, the endonuclease cleaves the concatemer into monomers, facilitating the monomerization. These monomers subsequently fold back into Nicking Loop™-converted circles, forming a nick in the stem that is ligated to complete circularization. This amplification process yields a new generation of CssDNA with opposite polarity to the original DNA strand, and the process can be repeated infinitely (Fig. 1B). With each generation, there is over 1000-fold increase in the number of circular molecules as quantified by qPCR.

The conversion of linear DNA by Nicking Loop™ is the key to infinite amplification. The efficiency of the conversion was determined by the abundance of the products detected via gel electrophoresis (Fig 1C). The incorporation of the Loop was highly efficient, representing 74% of all circular molecules. In the absence of the Loop, the linear DNA molecule is circularized, but the concatemeric amplification product of such molecule cannot be nicked and re-circularized, and is therefore not suitable for amplification cycles.

Each generation of the Nicking Loop™ amplification relies on coordinated action of nicking enzyme digestion of the amplified concatemer, and ligation to achieve monomer circularization, with the subsequent exonuclease degradation of any non-circular molecules to ensure purity of the product. The importance of these steps was demonstrated by gel electrophoresis showing the efficiency of the nicking enzyme in monomerization step (Fig. 1D), and the importance of ligation and subsequent exonuclease degradation to complete circularization (Supplementary Fig.1).

During monomerization, the amplified concatemer was effectively digested by a specific nicking enzyme into monomers (Fig. 1D). On the contrary, the concatemer remains intact if a nicking enzyme is absent or an incompatible recognition site is present, impairing the circularization. No product is synthetized in the absence of the template or polymerase and ligase enzymes during the Nicking Loop™ conversion, or in absence of amplification enzyme. However, if Loop is left out during Nicking Loop™ conversion, the non-converted circular molecules may amplify but monomerization and subsequent circularization did not occur, and any leftover DNA is degraded by exonucleases.

After successful monomerization, each monomer is simultaneously folded back into the Nicking Loop™-converted structure, forming a nick in the Loop stem that is ligated, and the circularization is completed. The circularized molecules are resistant to exonuclease degradation, unlike the non-ligated monomer that remains linear and was completely digested (Supplementary Fig. 1).

### Nicking Loop™ amplification preserves the sequence composition of the sample DNA

Nicking Loop™ enables multiple amplification cycles with minimal distortion or amplification bias. To compare the amplification bias to PCR, Nicking Loop™-converted circles were prepared with decreasing variant allele fractions (VAFs) and amplified by three consecutive rounds of either Nicking Loop™ amplification or PCR (Fig 2).

**Figure 2.**
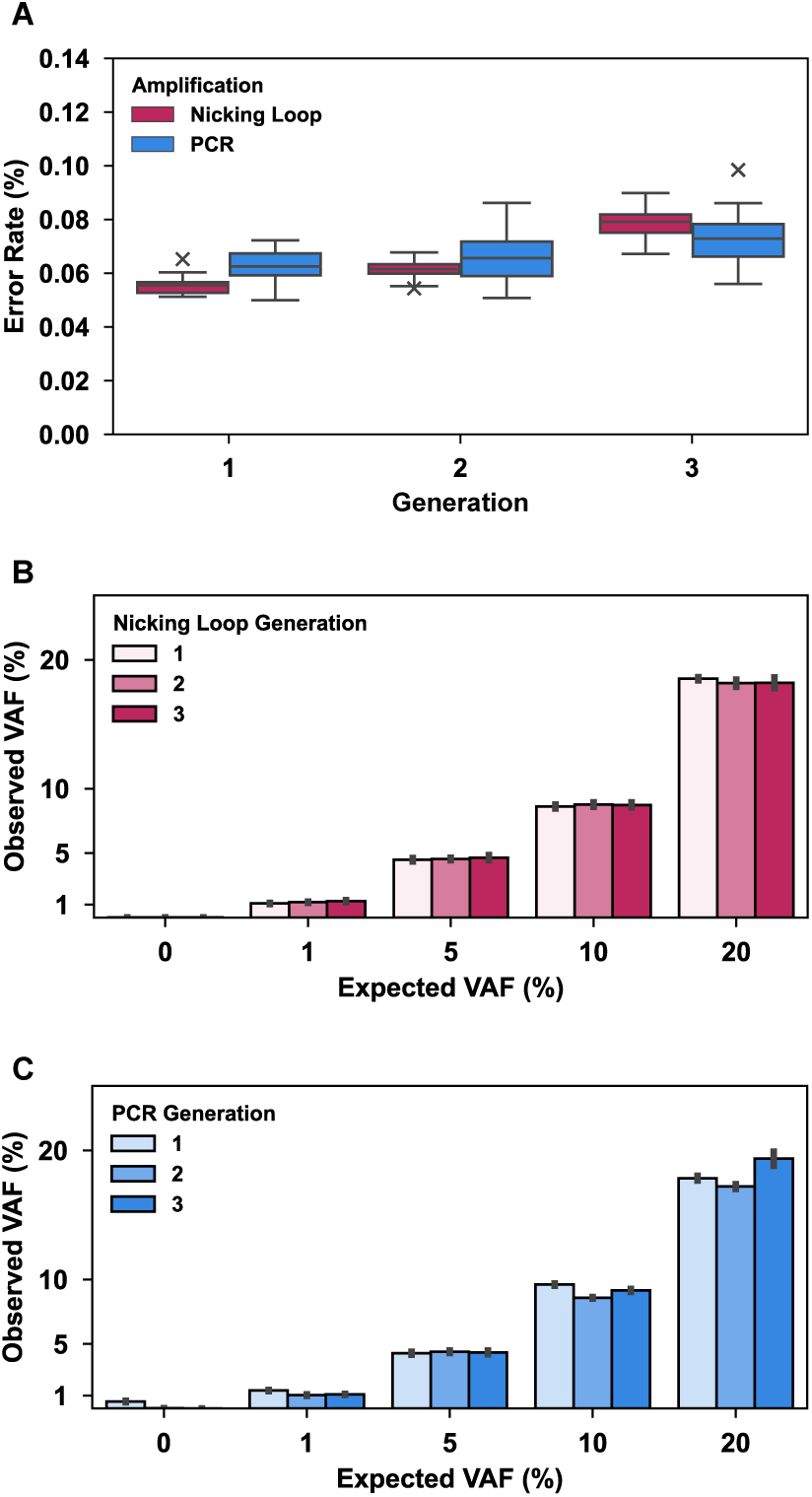
Amplification generations produced by Nicking Loop™ and PCR. **(A)** Error rate for Nicking Loop™ (pink) and PCR (blue). The error rate represents the proportion of mismatching nucleotides in the expected target gap nucleotides. The line in the boxplot represents the mean (instead of typical median) and error bars show standard deviation between 20 replicates. The variant allele frequency (VAF) variation between generations and sample replicas induced by **(B)** Nicking Loop™ and **(C)** PCR. The error bars represent standard deviation between 4 replicates.

The proportion of mismatching nucleotides relative to the expected target gap nucleotides was used as a proxy for the error rate (Fig. 2A). The base error rate comprises oligonucleotide synthesis errors, sequencing errors or errors caused by GapFill polymerase and is the shared baseline for both methods. The quantified difference between the error rates is hence caused solely by the amplification method. Both methods introduce new errors with each new generation. In the first two generations, the error rate of Nicking Loop™ (1^st^ generation: 0.056% ± 0.003%; 2^nd^ generation: 0.061% ± 0.004%), was lower than that of PCR (1^st^ generation: 0.063% ± 0.006%; 2^nd^ generation: 0.066% ± 0.010%). In the last generation, Nicking Loop™ (0.079% ± 0.006%) displayed higher error rates compared to PCR (0.073% ± 0.010%) but PCR consistently exhibited higher variation between the generations and replicas (Supplementary table S1).

To investigate how both amplification methods alter the template composition across generations, two pools of synthetic templates were mixed in different ratios to mimic different VAFs. Subsequently, the VAF variation across generations and replicates was determined by both methods (Fig. 2B; Fig. 2C). Overall, Nicking Loop™ preserved the VAF proportion across the generations, while PCR exhibited more variation. The detailed summary of observed VAFs across generations for Nicking Loop™ and PCR are available in Supplementary Table S2.

The most notable differences were observed at 1% VAF (Nicking Loop™: CV_1%_ = 6.04%, range: 1.09–1.27%; PCR: CV_1%_ = 13.36%, range: 1.04–1.39%), followed by 10% VAF (Nicking Loop™: CV_10%_ = 0.76%, range: 8.62–8.77%; PCR: CV_10%_ = 4.64%, range: 8.57–9.61%) and 20% VAF (Nicking Loop™: CV_20%_ = 0.85%, range: 18.19–18.53%; PCR: CV_20%_ = 4.99%, range: 17.20–19.36%). Contamination was observed in the first generation of PCR at 0% VAF but dropped out with subsequent generations. In PCR, similar trend could be observed at 1% VAF, probably caused by preferential amplification of most abundant templates, suppressing the amplification of the less abundant templates.

The VAF variation across generations for each probe in the panel is displayed in Figure 3. Mainly the probes 6 and 7, which were diluted tenfold relative to the rest of the panel were negatively influenced by PCR amplification. For the probe 6 at 1% VAF, PCR exhibited high variation across the generations (CV_1%_ = 104.07%, range: 0.00–1.63%) with no signal detected in 3^rd^ generation, while Nicking Loop™ exhibited consistent results (CV_1%_ = 9.79%, range: 0.98–1.07%). Similarly, for probe 7 in 3^rd^ PCR generation the signal dropped out at VAFs of 1%, 5% and 10% with high variation between generations (CV_1%_ = 92.84%, CV_5%_ = 70.96%, CV_10%_ = 81.14%). In this instance, the Nicking Loop™ preserved all the signal and maintained low variation between generations at VAFs of 1%, 5% and 10% (CV_1%_ = 20.35%, CV_5%_ = 8.03%, CV_10%_ = 12.64%). Overall, Nicking Loop™ exhibited less variation across generations and replicas for each probe (Supplementary Table S3). These results highlight the reproducibility of Nicking Loop™ amplification between different generations, particularly at low VAFs or low-abundance templates.

**Figure 3.**
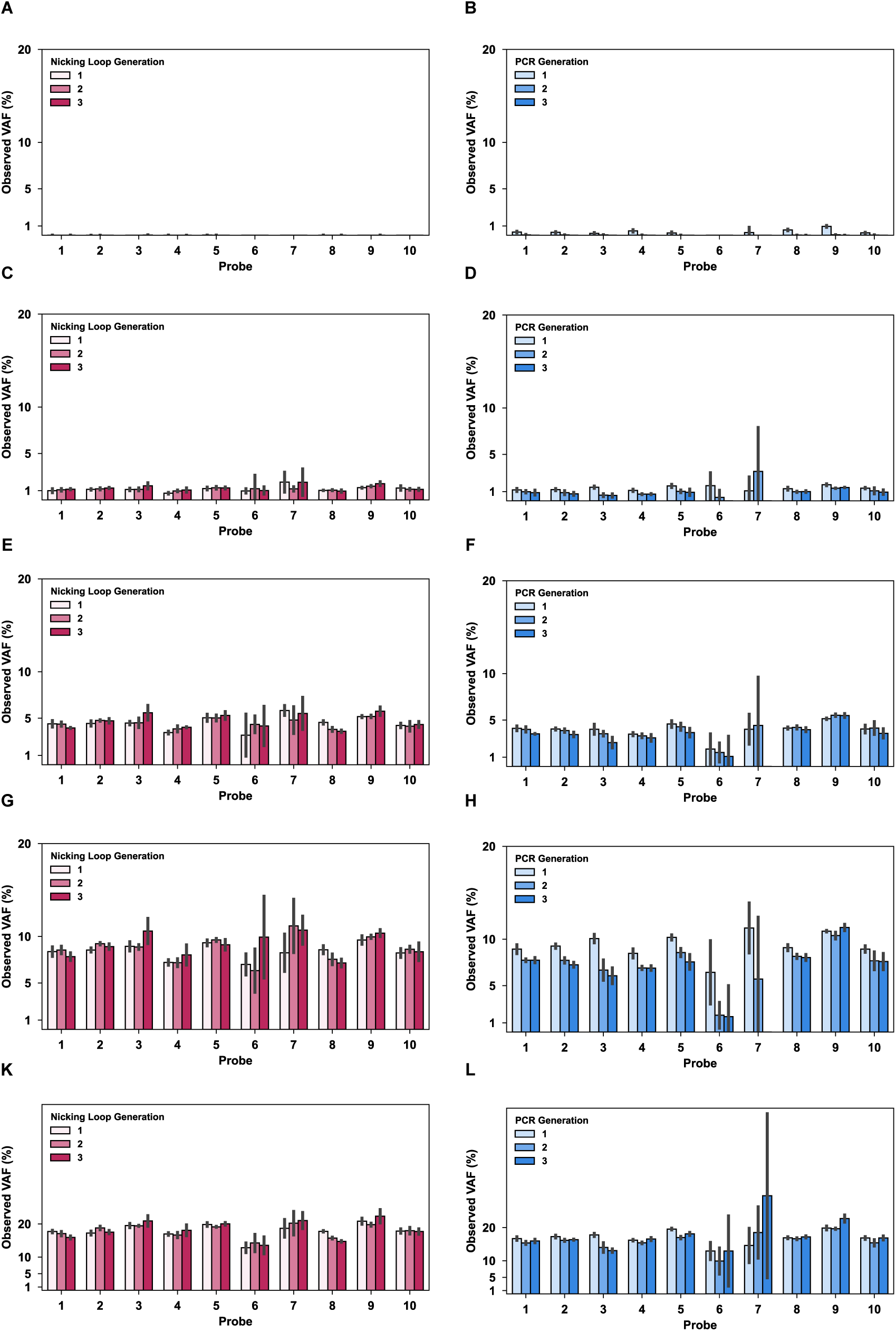
Performance of each probe across amplification generations by Nicking Loop™ and PCR. The generations produced by Nicking Loop™ (pink) corresponding to expected variant allele frequency (VAF) of **(A)**0%, **(C)** 1%, **(E)** 5%, **(G)** 10%, **(K)** 20%. The generations produced by PCR (blue) corresponding to expected VAF of **(B)** 0%, **(D)** 1%, **(F)** 5%, **(H)** 10%, **(L)** 20%. The error bars represent standard deviation between 4 replicates.

### Nicking Loop™ permits early sample indexing

The loop-region can carry the sample indices as well as UMIs, enabling early sample pooling and error correction. Therefore, it is essential that these nucleotides are customizable and do not hinder any aspects of the reaction. To assess the impact of the sequence variability within the loop-region, the Loops were tested in competition assay, contrary to the typical setup with single oligonucleotide in the reaction. An equimolar mixture of 25 oligonucleotides with varying centremost nucleotides (Supplementary table S4) was subjected to the usual workflow in ten replicas. The performance of each oligonucleotide was evaluated based on their proportion of total on-target reads obtained from the reaction (Fig. 4A).

**Figure 4.**
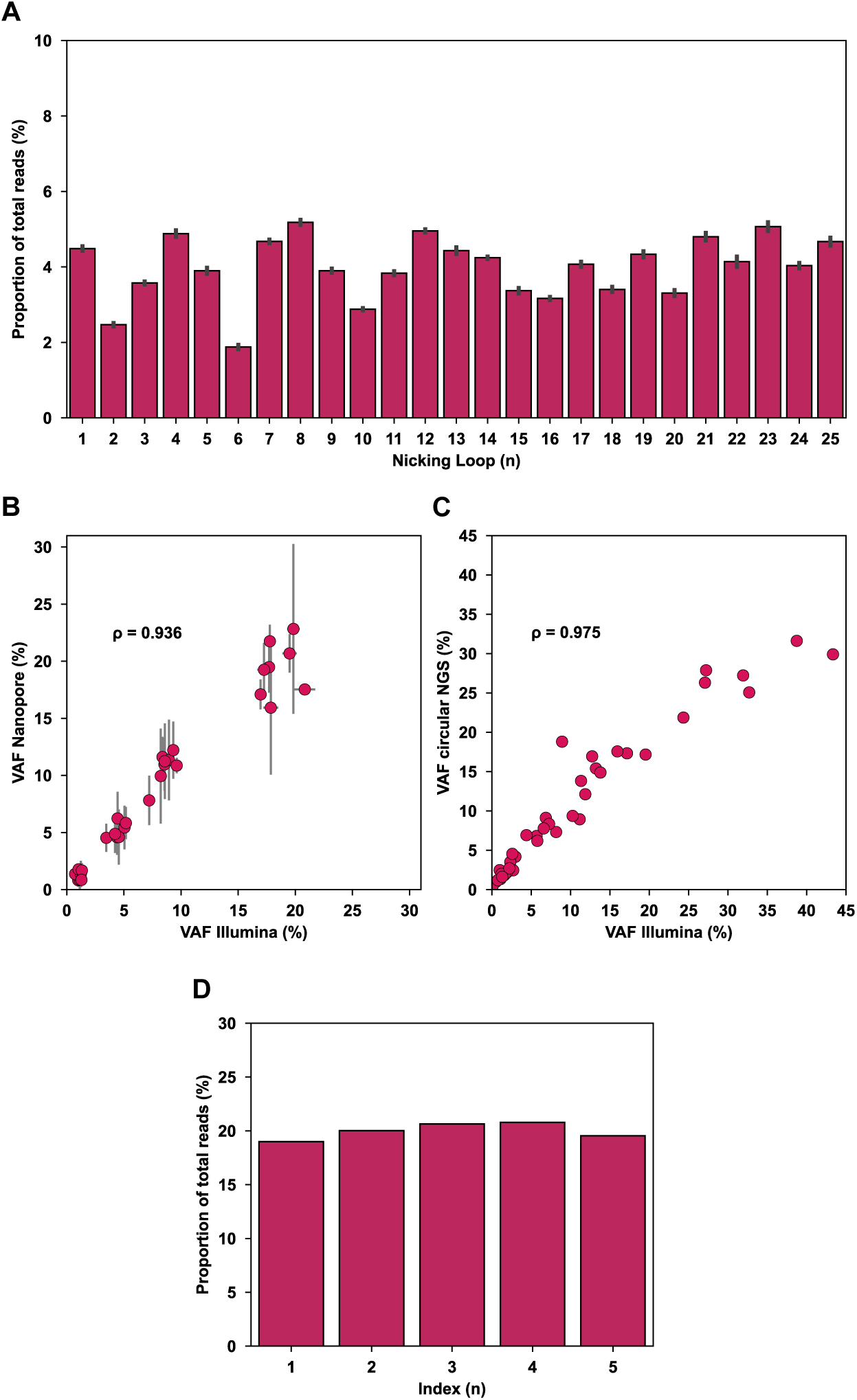
The application of Nicking Loop™. **(A)** Incorporation efficiency of the Loop. Twenty-five distinct Loops with varying sequences in the loop-region competed for their incorporation during the circular single-stranded DNA conversion. Each bar represents a proportion of total on-target reads (%) the given Loop produced. Error bars show standard deviation between 10 replicates. **(B)** Nicking Loop™ compatibility across next-generation sequencing (NGS) platforms – **(B)** Oxford Nanopore and **(C)** circular NGS - with Illumina MiSeq as reference. Different Nicking Loop™ intermediates were used as a library – amplified concatemer for Oxford Nanopore, Nicking Loop™-converted circles for circular NGS and linear template for Illumina MiSeq. The variant allele frequencies (%) detected in libraries show a strong correlation (Spearman’s rank correlation coefficient) for Oxford Nanopore (ρ = 0.936) and circular NGS (ρ = 0.975) referred to linear Illumina MiSeq libraries. **(D)** Proportion of total on-target reads (%) for each unique sample index (n) encoded in the Loop for direct circular sequencing, demonstrating efficient sample multiplexing.

The results show comparable performance between most oligonucleotides without excessive enrichment, suggesting that loop-indexing approach is compatible with pooling of multiple samples. Oligonucleotides 21–25 mimicked the presence of UMIs and exhibited minimal differences in performance. There was no clear relation of varying performance to GC content, or the number of nucleotides in the oligonucleotide sequence. Lastly, the variation between 10 replicas was low, demonstrating robustness of the reaction.

### Nicking Loop™ is compatible with read-out on various NGS platforms

Nicking Loop™ enables a versatile library preparation compatible with widely used NGS platforms regardless of their platform-specific chemistry. To demonstrate the compatibility of Nicking Loop™ with different sequencing platforms, the Nicking Loop™ concatemers were sequenced using Oxford Nanopore (Fig. 4B), Nicking Loop™-converted circles were sequenced using a direct circular sequencing platform (Fig. 4C), and linear Illumina MiSeq library served as the reference. The linear Illumina MiSeq library was prepared from the matching circular templates of the compared platform. Despite different intermediates used for each platform, sequencing results revealed strong agreement of VAFs for Oxford Nanopore (ρ = 0.936) and circular NGS (ρ = 0.975) with the Illumina MiSeq reference, underscoring the compatibility of Nicking Loop™ with multiple sequencing platforms.

The circular templates were engineered to include the necessary flow cell binding motifs, all required primer binding sites for circular NGS platform and unique sample indices incorporated into the Loop. Furthermore, Figure 4D illustrates clear separation of distinct samples based on the Loop-encoded sample indices, highlighting efficient sample demultiplexing from circular templates. Collectively, these findings confirm that Nicking Loop™ is not only compatible with various NGS platforms but also enables efficient transition towards direct CssDNA sequencing. Moreover, the ability to accurately identify samples via Loop-indices provides demonstration of early indexing of circular library structures that enhances sequencing performance and streamlines library preparation workflows.

## DISCUSSION

Nicking Loop™ is a novel approach for permitting conversion of ssDNA or dsDNA into structurally defined CssDNA molecules, enabling repeated high-fidelity linear DNA amplification, and facilitating direct or near-direct sequencing on circular or nanopore sequencing platforms, respectively. The amplification can be precisely controlled by adjusting the number of amplification generations, thus fine-tuning the DNA yield and strand polarity. Even multiple amplification cycles preserve the original information content with high accuracy, showing promise for highly challenging applications such as minimal residual disease detection, while streamlining basic assay preparation steps. Furthermore, Nicking Loop™ provides a versatile and powerful tool for DNA data storage applications by minimizing amplification bias, preserving original template composition and enhancing overall copying and sequencing accuracy.

In this study, Nicking Loop™ amplification and PCR were systematically compared for respective error rates and VAF distortion. Nicking Loop™ amplification demonstrated partially lower error accumulation across generations and reduced variability between replicate experiments compared to PCR. The read-out of most Nicking Loop™ experiments was conducted on Illumina platforms, which necessitates the incorporation of flow cell binding sequences via PCR – a step that inherently introduces PCR-associated errors. Consequently, the full benefits of Nicking Loop™ amplification may not have been completely realized under these conditions. It is anticipated that with the adoption of circular NGS platforms, where PCR amplification steps can be omitted, the differences in error rates between PCR and Nicking Loop™ amplification would become even more pronounced.

Nicking Loop™ accurately preserved the original VAFs and relative proportion of low-abundance templates. Conversely, distortions at low VAFs and low-abundance templates were evident with PCR, aligning well with reported findings (12, 13). Therefore, Nicking Loop™ shows potential for very sensitive detection applications, e.g. the detection of low-abundance targets amidst high-abundance DNA background in oncological applications. Additionally, the low-abundance templates can be selectively amplified with Nicking Loop™ simply using template-specific amplification primers (unpublished data).

This study evaluated how sequence variability in the Loop might affect its ability to incorporate into the Nicking Loop™-converted circles. No preferential enrichment of any specific sequence motifs within the Loops was observed, and majority of the Loops performed comparably. These findings suggest that Nicking Loop™ can support early sample pooling strategies and can carry UMIs without compromising the amplification or sequencing performance. This has the potential to significantly streamline NGS library preparation workflows and reduce the overall cost of library preparation and sequencing.

The Nicking Loop™-converted circular molecules serve as ready NGS libraries directly compatible with sequencing platforms utilizing circular DNA molecules as sequencing templates such as Element Bioscience’s AVITI (4), Pacific Bioscience’s Onso (5), and MGI’s DNBSEQ (6). Direct use of circular DNA libraries reduces workflow steps, enabling rapid and simplified NGS workflows. The long concatemeric DNA molecules generated by RCA serve as templates for Oxford Nanopore (7), expanding the applicability of the method across different sequencing platforms. Moreover, nicked linear monomeric amplification products can be sequenced using platforms designed for linear library molecules such as Illumina or Ion Torrent.

Any products of the Nicking Loop™ process can be subjected to further amplification to achieve required library concentration, thus mitigating the challenges associated with low input material. Preliminary evaluations of various sequencing platforms utilizing Nicking Loop™-derived templates have shown promising results, further highlighting the method’s versatility and potential for broad adoption.

Besides the sequencing applications, the stability of the CssDNA libraries over linear DNA libraries points towards utility in prolonged ambient storage of DNA samples (unpublished data). This, coupled with the highly accurate replication of information, including low-abundance targets, points towards applications within the DNA data storage field.

The Nicking Loop™ is poised to lead a paradigm shift in how circular DNA is prepared, replicated and sequenced, moving away from rigid linear library structure to more versatile circular molecules. This permits improvements in NGS library preparation, offering benefits in speed, cost-efficiency and workflow simplicity. The low infrastructural needs of Nicking Loop™ have the potential to unlock NGS and DNA data storage applications even in non-laboratory settings.

## DATA AVAILABILITY

The data underlying this pre-print will be shared on reasonable request to the corresponding author.

## SUPPLEMENTARY DATA

Supplementary Data are available online.

## AUTHOR CONTRIBUTIONS

Simona Adamusová: Conceptualization, Formal analysis, Investigation, Methodology, Visualization, Writing—original draft.

Nea Laine: Conceptualization, Data Curation, Formal analysis, Methodology, Software, Visualization, Writing—original draft.

Anttoni Korkiakoski: Data curation, Formal analysis, Methodology, Software, Writing—review & editing.

Tatu Hirvonen: Investigation, Methodology, Writing—review & editing.

Anna Musku: Investigation, Writing—review & editing.

Tuula Rantasalo: Investigation, Writing—review & editing.

Jorma Kim: Data curation, Writing—review & editing.

Juuso Blomster: Funding, Writing—review & editing.

Jukka Laine: Writing—review & editing.

Manu Tamminen: Conceptualization, Funding acquisition, Methodology, Project administration, Supervision, Writing—review & editing.

Juha-Pekka Pursiheimo: Conceptualization, Investigation, Methodology, Project administration, Supervision, Writing—original draft.

## ACKNOWLEDGEMENTS

The authors would like to express their gratitude to Voima Ventures, Avohoidon Tutkimussäätiö and Almaral for support and funding.

## ABBREVIATIONS

CssDNA: Circular single-stranded DNA
dsDNA: Double-stranded DNA
qPCR: Quantitative real-time PCR
RCA: Rolling circle amplification
ssDNA: Single-stranded DNA
UMI: Unique molecular identifier
VAF: Variant allele frequency

