## Supplementary Material for "Nicking Loop™: Streamlined circular DNA libraries for precision genomics, DNA data storage and universal NGS read-out"

**Supplementary Figure 1.** The circularization step depicted by gel electrophoresis (Tapestation – HS RNA ScreenTape). The folded monomers in absence of the ligase remain in the linear ssDNA form and after addition of ligase the monomers circularized. After the circularization is completed, the linear ssDNA is degraded by exonucleases, while circularized molecules resist degradation and remains intact.

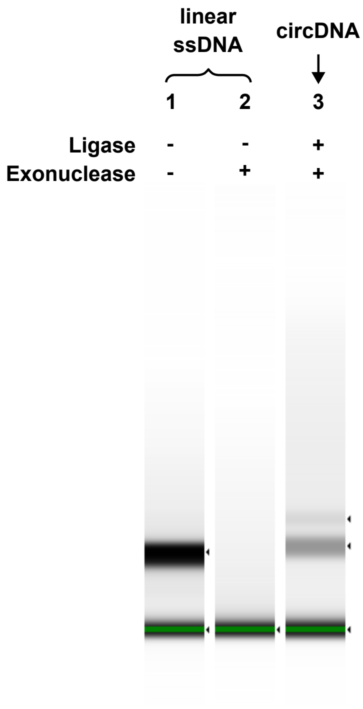

**Supplementary Table S1.** Error rate values (%) ± replicate standard deviation (%) for each generation of Nicking Loop™ and PCR.

| Method | Generation | Error rate (%) |
|---|---|---|
| Nicking Loop™ | 1 | 0.056 ± 0.003 |
| Nicking Loop™ | 2 | 0.061 ± 0.004 |
| Nicking Loop™ | 3 | 0.079 ± 0.006 |
| PCR | 1 | 0.063 ± 0.006 |
| PCR | 2 | 0.066 ± 0.010 |
| PCR | 3 | 0.073 ± 0.010 |

**Supplementary Table S2.** Summary of observed VAFs across generations for Nicking Loop™ and PCR. The table includes expected VAF (%), mean VAF (%) with replicate standard deviation (STD) VAF (%) and coefficient of variation (CV) across generations for given method and expected VAF.

| Expected VAF (%) | Method | Generation | Mean VAF (%) | STD VAF (%) | CV (%) |
|---|---|---|---|---|---|
| 0 | Nicking Loop™ | 1 | 0.00 | 0.00 | 44.09 |
| | Nicking Loop™ | 2 | 0.00 | 0.00 | |
| | Nicking Loop™ | 3 | 0.00 | 0.00 | |
| | PCR | 1 | 0.54 | 0.06 | 130.56 |
| | PCR | 2 | 0.03 | 0.01 | |
| | PCR | 3 | 0.00 | 0.00 | |
| 1 | Nicking Loop™ | 1 | 1.09 | 0.05 | 6.04 |
| | Nicking Loop™ | 2 | 1.19 | 0.04 | |
| | Nicking Loop™ | 3 | 1.27 | 0.09 | |
| | PCR | 1 | 1.39 | 0.05 | 13.36 |
| | PCR | 2 | 1.04 | 0.05 | |
| | PCR | 3 | 1.09 | 0.04 | |
| 5 | Nicking Loop™ | 1 | 4.48 | 0.14 | 1.47 |
| | Nicking Loop™ | 2 | 4.54 | 0.10 | |
| | Nicking Loop™ | 3 | 4.65 | 0.20 | |
| | PCR | 1 | 4.29 | 0.11 | 1.11 |
| | PCR | 2 | 4.40 | 0.09 | |
| | PCR | 3 | 4.35 | 0.15 | |
| 10 | Nicking Loop™ | 1 | 8.62 | 0.15 | 0.76 |
| | Nicking Loop™ | 2 | 8.77 | 0.17 | |
| | Nicking Loop™ | 3 | 8.74 | 0.20 | |
| | PCR | 1 | 9.61 | 0.09 | 4.64 |
| | PCR | 2 | 8.57 | 0.06 | |
| | PCR | 3 | 9.15 | 0.12 | |
| 20 | Nicking Loop™ | 1 | 18.53 | 0.13 | 0.85 |
| | Nicking Loop™ | 2 | 18.19 | 0.28 | |
| | Nicking Loop™ | 3 | 18.21 | 0.41 | |
| | PCR | 1 | 17.85 | 0.18 | 4.99 |
| | PCR | 2 | 17.20 | 0.18 | |
| | PCR | 3 | 19.36 | 0.54 | |

**Supplementary Table S3**. Provided as separate file.
Summary of observed VAFs for each probe across generations for Nicking Loop™ and PCR. Each sheet represents expected VAF (%) and provides an overview of mean VAF (%) with replicate standard deviation (STD) VAF (%) and coefficient of variation (CV) for each probe across generations for given method.

**Supplementary Table S4.** Structural differences of Loop and proportion of total reads (%) each oligomer captured in the sequencing (on-target reads only).

| Loop | Centremost nucleotides of loop-region | GC content (%) | Proportion of total reads (%) | Proportion of total reads STD (%) |
|---|---|---|---|---|
| 1 | TTCTGCCT | 50 | 4.49 | 0.06 |
| 2 | GCTCAGGA | 62.5 | 2.47 | 0.05 |
| 3 | AGGAGTCC | 62.5 | 3.58 | 0.04 |
| 4 | CATGCCTA | 50 | 4.88 | 0.09 |
| 5 | GTAGAGAG | 50 | 3.90 | 0.08 |
| 6 | CCTCTCTG | 62.5 | 1.88 | 0.06 |
| 7 | TGCCTCTT | 50 | 4.68 | 0.05 |
| 8 | TAGCGAGT | 50 | 5.18 | 0.07 |
| 9 | CCTGAGAT | 50 | 3.90 | 0.05 |
| 10 | GTAGCTCC | 62.5 | 2.88 | 0.03 |
| 11 | AGGCTCCG | 75 | 3.83 | 0.06 |
| 12 | GCAGCGTA | 62.5 | 4.96 | 0.04 |
| 13 | CTGCGCAT | 62.5 | 4.43 | 0.09 |
| 14 | GAGCGCTA | 62.5 | 4.24 | 0.03 |
| 15 | CGCTCAGT | 62.5 | 3.37 | 0.07 |
| 16 | GTCTTAGG | 50 | 3.17 | 0.04 |
| 17 | ACTGATCG | 50 | 4.07 | 0.07 |
| 18 | TAGCTGCA | 50 | 3.40 | 0.07 |
| 19 | GACGTCGA | 62.5 | 4.34 | 0.08 |
| 20 | CGTCTGAC | 62.5 | 3.31 | 0.08 |
| 21 | NNTGNNTCANN | N/A | 4.80 | 0.10 |
| 22 | NNNTGNNTANNN | N/A | 4.14 | 0.14 |
| 23 | NNNNNNNN | N/A | 5.07 | 0.12 |
| 24 | NNNNNNNNNN | N/A | 4.03 | 0.07 |
| 25 | NNNNNNNNNNNN | N/A | 4.67 | 0.11 |
